# Ability to monitor deviations of own movement without visual feedback

**DOI:** 10.1101/2024.05.09.593337

**Authors:** I.M. Brandt, S.M. Bjerregaard, D.B. Jensen, M.G. Giessing, T.G. Grünbaum, M.S. Christensen

## Abstract

How conscious sensations of movement relates to signals essential for movement control remains under investigation. This question is typically investigated using visuomotor tasks, in which sensation of movement is disturbed by conflicting visual feedback. The present study uses metacognitive judgements to investigate conscious access to movement signals, unchallenged by visual signals, in an index finger force task. We found that some participants can correctly assign metacognitive judgements (MCJs) to their own force, suggesting that participants do indeed have metacognitive access to sensorimotor signals. We found no correlation between metacognitive access to sensorimotor signal and variance in motor performance. Further, we found that in this purely sensorimotor task, internal focus of attention reduces variability in force compared to external focus of attention. Our results indicate that not only is it possible to access sensorimotor information, it is also possible to use focus of attention to reduce variability in force performance.

## Introduction

Movement is controlled through a complex interaction between the brain, the spinal cord, and efferent and afferent signals. The brain and the spinal cord process and integrate these signals to coordinate and execute precise movements of our limbs. Moving one’s body *feels like something*. The relationship between the conscious sensations of movement and the signals essential for movement control and coordination is still under investigation. In this study we examined conscious access to own body movements with metacognitive judgments in an isometric index finger force task without visual feedback. We also manipulated focus of attention (FOA) to investigate the effect of conscious focus on precision in movements.

It is an ongoing scientific discussion to which extend we have conscious access to sensorimotor signals, and which of the underlying signals this sensation relies on. Different lines of research point in different directions. First, there are studies investigating whether manipulating signals separately (cortical-, motor-, and sensory signals) gives rise to a sensation. These studies generally find that all signals can give rise to a sensation of movement (Christensen et al., 2010; Gandevia et al., 2006; McCloskey, 1973). Second, there are studies that introduce a discrepancy between visual and proprioceptive feedback. These studies generally find that we tend to rely on visual feedback, and that even large deviations in proprioceptive feedback goes unnoticed. These studies suggest that while such information may be crucial for movement control, our conscious awareness of the signals might not be essential (Fourneret and Jeannerod, 1998; Maselli et al., 2023). The last line of research has taken an important step by developing metacognitive tasks. Metacognitive ability refers to the capacity to monitor one’s own cognitive processes (Arbuzova et al., 2021), and in these tasks, participants report their subjective experience of the movement in each trial. Using this method, a metacognitive study by Pereira et al. (2023) builds upon the visuomotor perturbation studies and, similarly, find a lack of ability to report perturbations (Pereira et al., 2023). They further found that although participants do not report to have noticed perturbations in their movements, metacognitive confidence judgments distinguish between trials with and without perturbations (Pereira et al., 2023). So, it can be argued that the motor signals inform their subjective confidence in the movement assessment, and are thus to some degree accessible.

Metacognitive assessments have to a high degree been used to study visual perception and to a lesser degree to investigate movement perception. The studies investigating metacognition of movement build on tasks that have a variation of visual and proprioceptive feedback. In an elegant study by Charles et al. (2020), the authors investigated the relative contribution of efferent and afferent signals in motor awareness and metacognitive judgments of the movement by comparing metacognitive judgments between three conditions: an active and a passive condition of a finger flexion/extension movement, and a visual condition (Charles et al., 2020). They found higher accuracy in judging whether a visual probe appeared ahead of or behind their finger in the active compared to the passive and visual condition, even when using a staircasing method to equate performance between conditions. The staircasing method made the gap between the probe and the finger position smaller in the active condition. These results indicate that sensorimotor signals contribute to the decision accuracy of the position of the probe relative to the hand. They attribute this heightened accuracy to access to voluntary motor commands and proprioceptive information (Charles et al., 2020). They used a second-order signal detection theory method to control for first-order accuracy and confidence bias when assessing metacognitive ability, and they found no significant differences in metacognitive efficiency between the conditions (Charles et al., 2020). These findings indicate that sensorimotor signals contribute to the accuracy of the conscious experience of the position of the probe, but do not inform metacognitive efficiency.

Arbuzova et al. (2021) designed two experiments with three conditions each: 1) a motor, 2) a visuomotor, and 3) a visual ball throwing condition (Arbuzova et al., 2021). In the first experiment they had to choose the correct trajectory between two trajectories, and then rate the confidence in their decision. In the second experiment, they were to choose between two angles, and rate the confidence in their decision. They found that the accuracy for choosing the correct trajectory in both experiments were higher for visuomotor and the visual condition compared to the motor condition. However, they found similar metacognitive efficiency across conditions.

These studies include visual components in the tasks. It is well-known that visual information dominates over proprioception (Ernst and Banks, 2002; Rock and Victor, 1964). Judging one’s movement based on visual cues in the external world relies on mental transformations between the executed movement and corresponding visual cues. To avoid the effect of visual feedback and specifically assess our ability to consciously access motor signals, we have composed an experimental setup of an index finger force task without a visual component.

The present study differs from prior metacognitive studies of own body movement in two main ways. First, prior metacognitive studies of movement are engaged with the question of metacognitive efficiency and sensitivity, questions of discriminating between own correct and incorrect responses in a forced choice task. In this paper we use metacognitive judgments to investigate the access to sensorimotor signals, not the ability to evaluate own judgments. We will call this metacognitive access. Therefore, we did not create a task that allowed us to use the second-order signal detection theory method that could give us an estimate of the metacognitive sensitivity. The downside of this is that the ability to assess deviations in one’s own movement is affected by the ability to perform the movement. We accommodate this by comparing metacognitive ability to variability in movement performance. The strength is that we investigate conscious access to movement signals, rather than the ability to evaluate own judgments. Second, we have excluded visual feedback from our setup to evaluate conscious access to sensorimotor signals unaffected by visual information.

Another way to investigate the role of conscious awareness in control of movement is through investigating whether an internal focus of attention on the feeling of the movement influences the precision of the movement. The literature comparing external focus of attention (EFOA) to internal focus of attention (IFOA) has generally found an advantage of EFOA (Lohse et al., 2014b; Wulf, 2013). However, in these studies the movement outcome is typically to accomplish an effect in the external world, such as throwing arrows at a dart board, which is guided by vision (Lohse et al., 2014b). This naturally gives a disadvantage to IFOA, in which participants are instructed to draw their attention away from the visual information. In this paper we investigate whether EFOA or IFOA provides an advantage to movement precision when there is no informative visual information giving an advantage to the EFOA condition. We hypothesize that the IFOA condition allows participants to be more precise in their movements in the absence of visual signals.

We aim to assess if agents have accurate conscious access to movement signals in situations where movement signals are unchallenged by visual signals. We want to use metacognitive judgments to assess conscious access to movement signals rather than the ability to evaluate one’s own judgments. Given studies manipulating afferent and efferent signals and metacognitive studies of reaching movements, our hypothesis is that participants do in fact have (metacognitive) access to deviations in force in an index finger motor task without visual feedback.

## Methods

### Participants

Thirty healthy participants completed the study (21 female, mean age 24.2, age range 18-35). We recruited participants through forsøgsperson.dk and University of Copenhagen. We obtained written informed consent from all participants for their participation and data collection. The study was conducted in compliance with the declaration of Helsinki and data processing adhered to the EU General Data Protection Regulation and guidelines were approved by UCPH (2004334 – 4242). The protocol was approved by The Capital Region of Denmark (H-21061035). Participants received compensation at a rate of 150 DDK per hour. 3 participants were borderline ambidextrous (laterality quotient = 40), the remaining were right-handed when assessed with the Edinburgh handedness inventory (Oldfield, 1971) (71 +/-19.9 (mean +/-1 sd)). Additional questionnaires not relevant to this study were also administered.

### Experimental Design

This study investigated the ability to consciously access movement signals in a task without visual feedback. We did this with two approaches, one in which we instructed the participants to report a metacognitive judgment of their movement, and one in which we asked participants to hold their focus of attention on either internal or external signals.

Participants were seated at a table with both arms resting horizontally. The right forearm was positioned semi-prone, with the palm facing medially and the index finger extended straight. Participants were instructed to press with their index finger on a stationary force transducer with one of three force levels, 1, 3, or 5, on a scale from 1-10 where 10 correspond to maximal voluntary contraction (MVC). The experiment was divided in three conditions; one in which they were instructed to report a metacognitive judgment for each trial, one in which they were instructed to hold an internal focus of attention (IFOA), and one in which they were instructed to hold an external focus of attention (EFOA). For each trial a tone instructed the force level they aimed for (1, 3, or 5) and obtained a measure of the maximum force level in that trial. Additionally, in the meta condition we obtained a metacognitive judgment for each trial. We randomized the order of condition between participants.

### Preparations

Participants completed the experiment during a single four-hour visit. We obtained informed consent and administered questionnaires at the outset. EEG measures and EMG measures were obtained, which were relevant for another study and will be reported elsewhere. Maximal voluntary contraction (MVC) of the right-hand index finger was measured with participants keeping their arm semi-prone, palm facing medially, and index finger extended straight. They exerted maximum force twice, each for three seconds, with a 90-second rest interval between attempts. The maximal force achieved during either attempt was recorded as their MVC. Visual feedback of the arm and hand was concealed by a cloth during all conditions. Breaks were encouraged and provided upon request.

### The task

Participants pressed their right index finger isometrically against a force transducer in a horizontal direction (Fig. 2). Their right-hand fingers were positioned straight ahead and supported by two sticks to provide lean support (Fig. 1). The task required participants to exert force at designated levels—1, 3, or 5—on a 1-10 scale, where 10 represents their maximum voluntary contraction (MVC). The scale thus corresponds to 10%, 30%, and 50% of their MVC, respectively. We refer to this scale as %MVC. To minimize fatigue, the highest level was set to five. Following the determination of their MVC, participants received verbal guidance on how to achieve force levels 1, 3, and 5, which were repeated 2-3 times. Subsequently, they had to rely solely on their sensorimotor perception, as no further external feedback was provided.

There are three conditions in the task, a meta condition, an internal focus of attention (IFOA) condition, and an external focus of attention (EFOA) condition. In the meta condition, for each trial, after they perform a force level, participants report metacognitive judgments of their motor performance. They report ‘under’, ‘approximate’ or ‘over’, using a computer keyboard, indicating how their press was compared to what they should have pressed. The participants were allowed to report multiple levels in case they were in doubt. These trials were excluded from the analysis. In the IFOA condition, participants did not report metacognitive judgements, but were instead instructed to focus on the ‘how it feels in the finger during the movement’. In the EFOA condition, participants were instructed to focus on an imagined apparatus resembling a thermometer, that provided visual feedback on the force level performed.

For all three conditions, participants were presented with three distinct auditory tones for each trial. The pitch of the initial tone indicated the target force level: low, middle, and high pitches corresponded to force levels 1, 3, and 5, respectively. After a variable inter-tone interval, the initial tone was followed by two tones in between which the movement should occur, the first was a ‘go-tone’, signaling the commencement of the action, and the second tone indicated that the movement should be finished. The second tone always followed the first tone with an 1.2 sec interval, which allowed the participants to learn to perform the force within this interval. Participants were instructed to exert the force rapidly in between the two consecutive tones but was not instructed to react fast. Participants completed six blocks of the meta condition and three blocks of the IFOA and EFOA conditions (except for the first two participants, who completed four blocks of each, and except for excluded blocks).

On average, each participant undertook 117 meta trials, 82.2 IFOA trials and 89.4 EFOA trials.

### Data acquisition

Spike2 (version 7.10, CED, Cambridge, UK) controlled the experimental setup including the auditory signals via a 1401 Micro Mk II analog-to-digital (AD) converter (Cambridge Electronic Design (CED), Cambridge, UK) and force signals. Data from the force transducer (UU2 load cell, DN-AM310 amplifier, Dacell, Seoul, Korea) were acquired at a sampling rate of 2000 Hz using the CED system. Data was securely stored on a protected drive. Surface EMG recordings were obtained from flexor digitorum profundus (FDP), first dorsal interosseous muscle (FDI), and extensor digitorum communis (EDC) and EEG data was collected using an active electrode EEG/EMG system (Active2, BioSemi, Amsterdam, The Netherlands) for purposes outside the scope of this paper.

### Data analysis and statistics

#### Force task performance

Data preprocessing was performed using the software MATLAB (version 9.13 (R2022b), The MathWorks Inc, Natick, MA, USA) and statistical analyses were performed using R (version 2022.12.0, https://www.r-project.org). Force was smoothed over 80 samples and baseline corrected in MATLAB. To identify whether force levels performed differed between aim levels, in R we initially defined a Bayesian linear mixed model for each condition (meta, IFOA, EFOA) with maximal force measure as the dependent variable and subject and aim (force level 1, 3, or 5) as independent variables. We used the ‘brms’ library in R (Bürkner, 2017) in R using STAN (Carpenter et al., 2017). Subsequently we used Bayesian hypothesis tests using the function ‘hypothesis’ from the ‘brms’ package to evaluate whether there was a difference between force aim levels.

#### Distance from force to %MVC

For each trial, we calculated the distance between the force level performed, and the %MVC level, as a measure of distance to instructed force level (DistanceToMVC). The %MVC level was calculated individually per participant from their respective MVC. DistanceToMVC was standardized per participant and per aim.

#### Distance from force to linear regression

To ensure that participants relied solely on sensorimotor signals, we did not provide feedback on their force levels during the task. As a results, their internal representation of the force level might not necessarily align with the instructed %MVC. To estimate each participant’s internal representation of the force level, we performed a linear regression per aim level for each block allowing drift over time. To measure the distance between the force performed in each trial and the internal representation, for each datapoint, we used a leave-one-out method to estimate the linear regression based on all other datapoints and predict the current datapoint. Subsequently, we calculated the distance between the performed force and the predicted force (DistanceToLM) for that trial.

#### Metacognitive judgments

To test whether there was a relationship between motor performance and MCJs, we evaluated the MCJs in relation to both DistanceToMVC and DistanceToLM. We did this by creating two Bayesian linear mixed models: one with DistanceToMVC and the other with DistanceToLM (both standardized per subject and aim) as the dependent variables. In both models, subject, MCJ, and aim were included as independent variables. Bayesian hypothesis tests were employed to evaluate whether DistanceToMVC and DistanceToLM differed between MCJs. Trials with multiple MCJs, indicating uncertainty, were excluded.

#### Testing the effect of condition

To test whether variance in motor performance differed between foci of attention, we used Bayesian hypothesis tests to evaluate whether DistanceToMVC and DistanceToLM differed between conditions meta, IFOA, and EFOA.

#### Relation between metacognitive ability and force performance

We tested whether there is a relationship between metacognitive ability and motor performance, in three distinct but related ways based on DistanceToLM. First, we determined percentage correct MCJs per participant. Correct MCJs were defined a positive DistanceToLM when MCJ was reported ‘above’ and negative DistanceToLM when MCJ was reported ‘below’. We used Bayesian mixed models and hypothesis tests to evaluate whether percentage correct was correlated with variance in motor performance, variance in DifferenceToLM. To account for a difference in the amount of reported ‘off trials’ (MCJ ‘below’ or ‘above’) compared to ‘on trials’ (MCJ ‘approximate’), we multiplied the percentage correct with the ratio ‘off trials’/all trials. Again, we used Bayesian mixed models and hypothesis tests to evaluate whether there was a correlation between (percentage correct*ratio of/all trials) and variance in DifferenceToLM. Lastly, we employed Bayesian mixed models and hypothesis tests to evaluate whether there was a correlation between a ratio of the DifferenceToLM variance in all trails and the DifferenceToLM variance in ‘approximate’ trials and variance in DistanceToLM.

## Results

We investigated whether participants were successful in performing the task. We evaluated whether they could produce three different force levels, without visual feedback. Figure 2 depictures the distributions of maximal force level for all participants, standardized per participant, under the three conditions, meta (Fig. 2a), IFOA (Fig. 2b), and EFOA (Fig. 2c).

**Figure 2.**
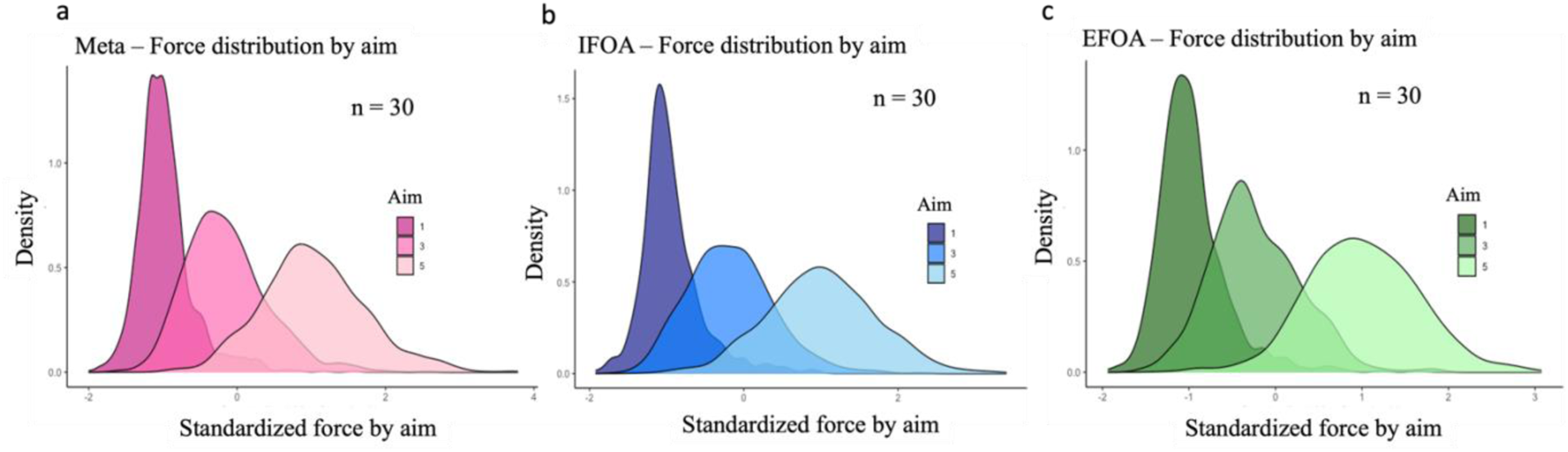
Distributions of maximal force levels by aim under the conditions a) Meta, b) Internal focus of attention (IFOA), and c) external focus of attention (EFOA). Data is normalized and standardized for each participant.

### Force distributed per aim level

Maximal force is distributed by level 1, 3, and 5. We estimated standardized maximal force as a function of subject and force level (Table 1). The estimate for maximal force increases with level across all conditions, and 95% credibility intervals (CI) do not overlap between force levels in any condition (Table 1), indicating that the force levels are distinguishable. An estimate of 0 corresponds to the individual mean across all trials for each participant. Bayesian hypothesis tests show that all force levels across each condition are different with infinite evidence ratio and a posterior probability of 1 (Table 1). Taken together, for each condition, participants can distinguish between the three different force levels.

**Table 1.**
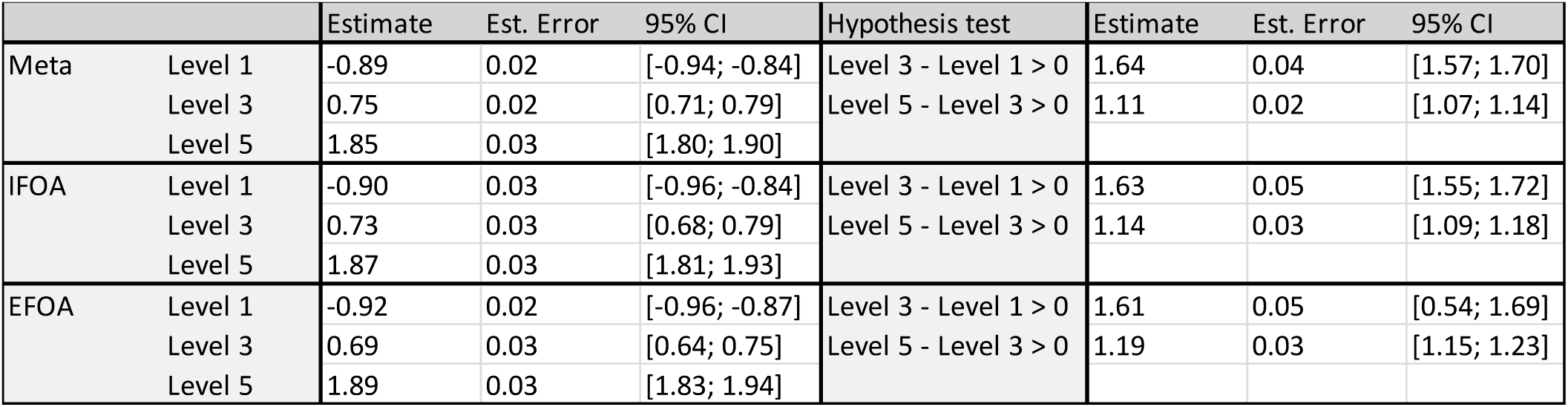
Distribution of maximal force by force level standardized per participant. An estimate of 0 corresponds to the mean force of the participant across all trials in the given condition. For all hypothesis tests across the three conditions, evidence ratios are infinite and posterior probabilities are 1. Data is normalized and standardized for each participant.

|  |  | Estimate | Est. Error | 95% CI | Hypothesis test | Estimate | Est. Error | 95% CI |
| --- | --- | --- | --- | --- | --- | --- | --- | --- |
| Meta | Level 1 | -0.89 | 0.02 | [-0.94; -0.84] | Level 3 - Level 1 > 0 | 1.64 | 0.04 | [1.57; 1.70] |
|  | Level 3 | 0.75 | 0.02 | [0.71; 0.79] | Level 5 - Level 3 > 0 | 1.11 | 0.02 | [1.07; 1.14] |
|  | Level 5 | 1.85 | 0.03 | [1.80; 1.90] |  |  |  |  |
| IFOA | Level 1 | -0.90 | 0.03 | [-0.96; -0.84] | Level 3 - Level 1 > 0 | 1.63 | 0.05 | [1.55; 1.72] |
|  | Level 3 | 0.73 | 0.03 | [0.68; 0.79] | Level 5 - Level 3 > 0 | 1.14 | 0.03 | [1.09; 1.18] |
|  | Level 5 | 1.87 | 0.03 | [1.81; 1.93] |  |  |  |  |
| EFOA | Level 1 | -0.92 | 0.02 | [-0.96; -0.87] | Level 3 - Level 1 > 0 | 1.61 | 0.05 | [0.54; 1.69] |
|  | Level 3 | 0.69 | 0.03 | [0.64; 0.75] | Level 5 - Level 3 > 0 | 1.19 | 0.03 | [1.15; 1.23] |
|  | Level 5 | 1.89 | 0.03 | [1.83; 1.94] |  |  |  |  |

We investigated whether condition (meta, IFOA, EFOA) influences the force distributions by evaluating whether there was a difference between the force distribution for each aim level (1,3,5) between conditions. We tested this with Bayesian hypothesis tests of the interaction between condition and aim level (Supplementary table 1). We found that the force distributions for each aim level did not differ between conditions, except for level 3, for which the estimate was larger in the meta condition compared to the EFOA condition (Est. = 0.06, Est. Error = 0.02 [0.02; 0.10], Evid. Ratio = 193, Post. Prob = 0.99) (Supplementary table 1). For no other aim level did the force distribution differ between conditions. These results suggest that the condition in general does not impact force level distributions.

### Metacognitive judgement and distance from force to %MVC

To evaluate whether the meta cognitive judgements reported by the participants are correct, first we determined the difference between the actual force produced and the force level they were asked to press on maximal force per trial normalized to individual MVC (not standardized) (Fig. 3). Second, we evaluated whether this difference is explanatory of the metacognitive judgement.

**Figure 3.**
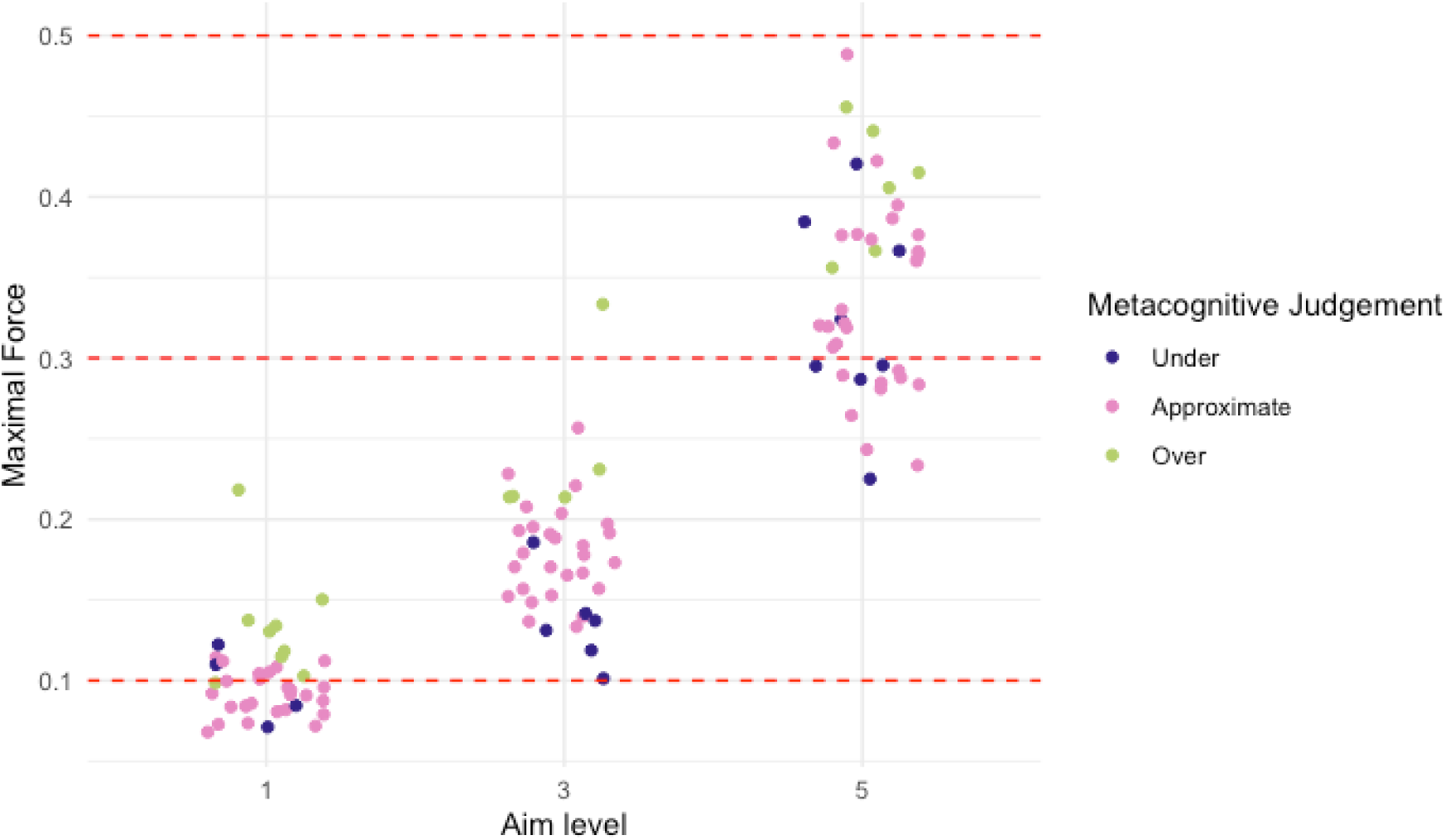
Force measurements for each level for a representative participant. Red dashed lines indicate the force level they should have performed (10% MVC, 30% MVC, 50% MVC). Each datapoint is colored by the metacognitive judgement (blue: below, pink: approximate, green: above). Data is normalized to individual MVC but not standardized. N = 119.

**Figure 4.**
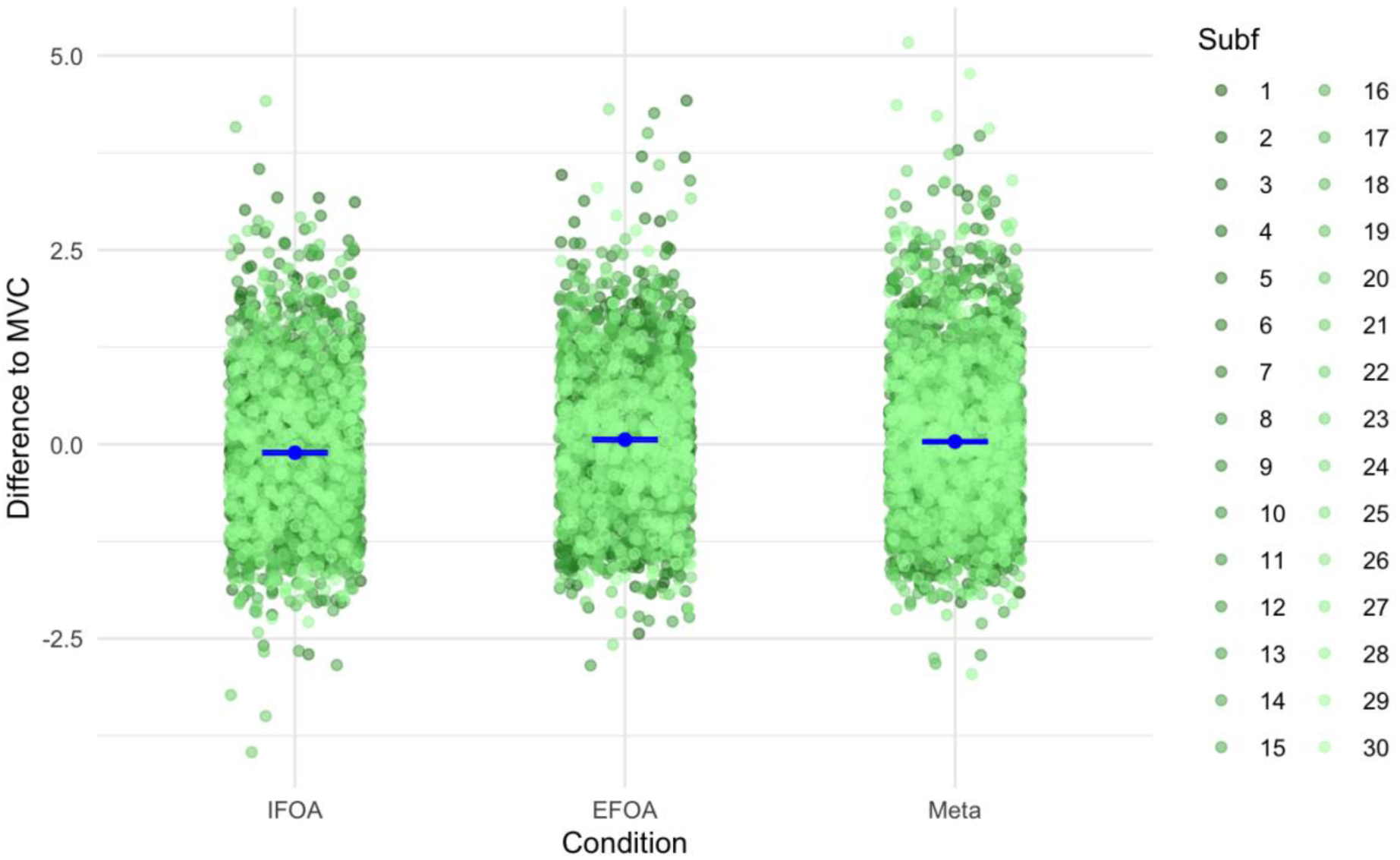
Difference between force and %MVC (10%, 30%, and 50% for aim level 1, 3, and 5, respectively) for conditions, meta, IFOA, and EFOA, standardized per subject and aim, colored by participant. 0 Signifies no difference, negative is undershooting and positive is overshooting. N = 8390.

To test whether condition has an impact on the difference between force and %MVC, we first standardized the difference to MVC by aim to align variance. Then we estimated a Bayesian linear regression model using brms, the difference between force and %MVC as a dependent variable of condition and subject. We found smaller estimates for the IFOA condition compared to both the meta and EFOA condition (Fig. 4, Table 2). This indicates that the participants are generally closer to the %MVC when focusing on internal signals compared to focusing on external signals or focusing on evaluating their performance.

**Table 2.** Left: Estimates of a brm model with distance between force and %MVC (DistanceToMVC) standardized per subject and aim as the dependent variable and condition and aim as the independent variables. Right: Hypothesis tests whether there is difference in the distance between force and %MVC between conditions. The distance to %MVC is lower under IFOA than both other conditions. This indicates that force is less variable around the internal representation of the force level in IFOA than in meta and EFOA.

|  | Estimate | Est. Error | 95% CI | Hypothesis test | Estimate | Est. Error | 95% CI | Evid. Ratio | Post. Prob |
| --- | --- | --- | --- | --- | --- | --- | --- | --- | --- |
| Intercept | -0.10 | 0.03 | [-0.16; -0.04] | Meta - IFOA > 0 | 0.25 | 0.05 | [0.17; 0.32] | Inf | 1.00 * |
| EFOA | 0.18 | 0.03 | [0.12; 0.23] | EFOA - IFOA > 0 | 0.27 | 0.05 | [0.20; 0.35] | Inf | 1.00 * |
| Meta | 0.15 | 0.02 | [0.10; 0.20] | EFOA - Meta > 0 | 0.03 | 0.02 | [-0.01; 0.07] | 6.93 | 0.87 |
| Level 3 | -0.00 | 0.03 | [-0.06; 0.05] |  |  |  |  |  |  |
| Level 5 | -0.04 | 0.03 | [-0.10; 0.01] |  |  |  |  |  |  |

To test whether the difference between force and MVC is explanatory of the metacognitive judgment in the meta condition, we estimated a Bayesian linear regression model with the difference between force and MVC as the dependent variable and metacognitive judgement and subject as the independent variables (Table 3). We performed Bayesian hypothesis tests to evaluate whether the estimates for the force levels differ between MCJ levels (Table 3, Fig. 5).

**Table 3.**
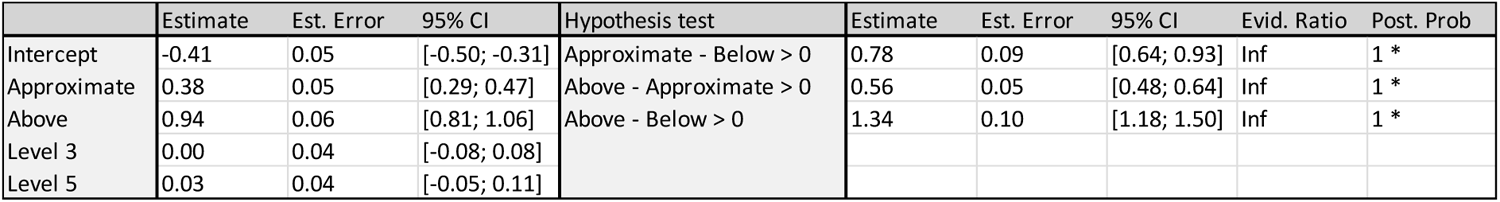
Estimates of a Bayesian linear regression model with distance to %MVC standardized by subject aim as the dependent variable and MCJ and aim as independent variables. Bayesian hypothesis tests test whether there is a difference between the MCJ levels.

**Figure 5.**
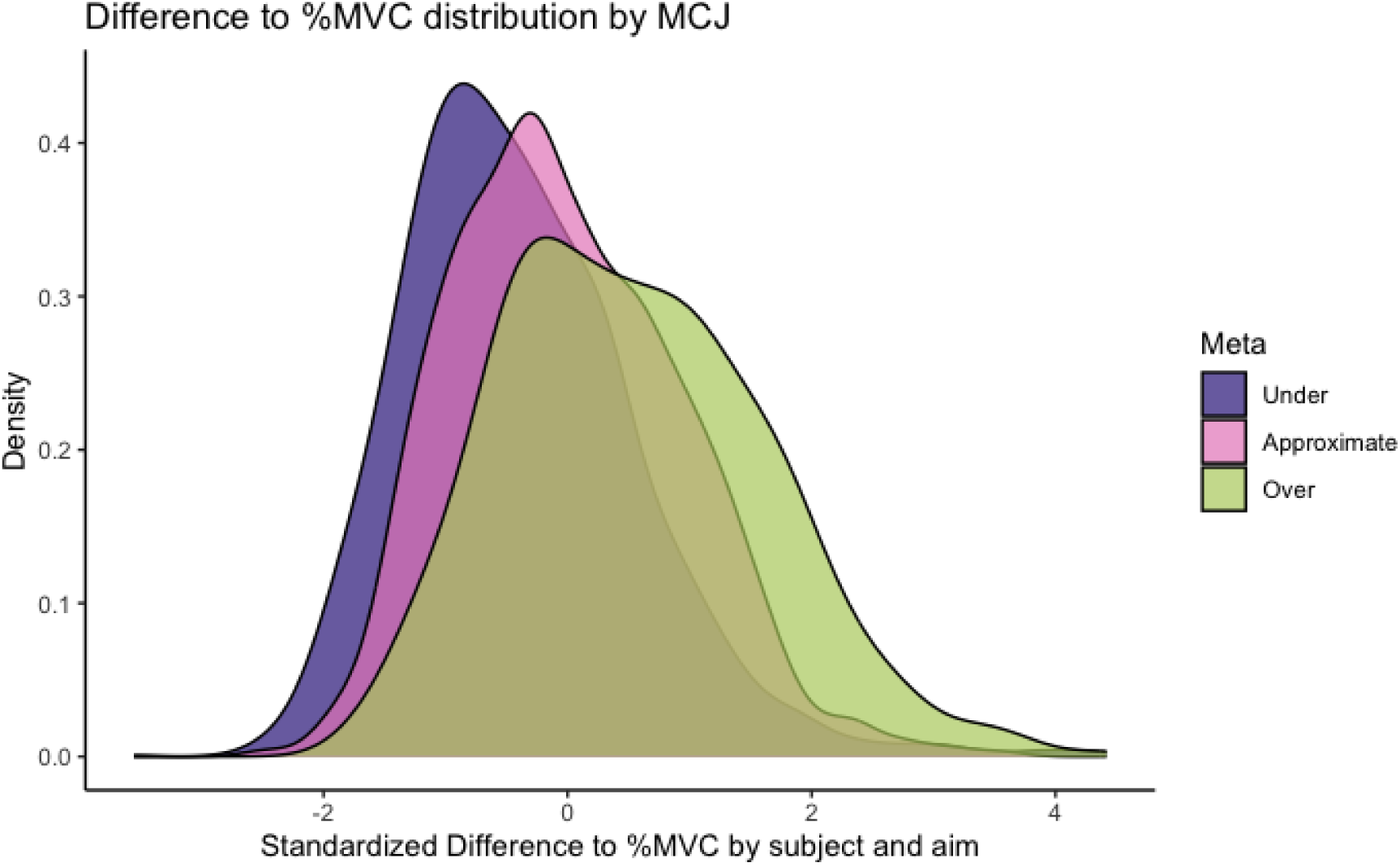
Distributions of distance to MVC standardized by participant and by aim, distributed by MCJ. N = 3510.

We find that the estimates of the distance to %MVC increase with MCJ, (Est.(below) = - 0.41, Est. Error = 0.0.5 [-0.50; -0.31], Est. (approximate) = 0.38, Est. Error = 0.05 [0.29; 0.47], Est (above) = 0.94, Est. Error = 0.06 [0.81; 1.06]). Hypothesis tests showed that these effects were significant with infinite evidence ratio and a posterior probability of 1 (Table 3). Together, the estimates of difference between force and %MVC increases with MCJ, indicating that participants correctly assign the MCJ.

### Meta cognitive judgement and distance from force to linear regression

Participants do not receive any visual input during the task, enabling the possibility of a drift in internal representation of the force levels over time. Further, the internal representations of the force levels are not necessarily centered around the right %MVC. To take this into account, we estimated linear regressions per force level for each participant and evaluated the difference between the actual force they produce, and the linear regression prediction using a leave-one-out analysis (Fig. 6).

**Figure 6.**
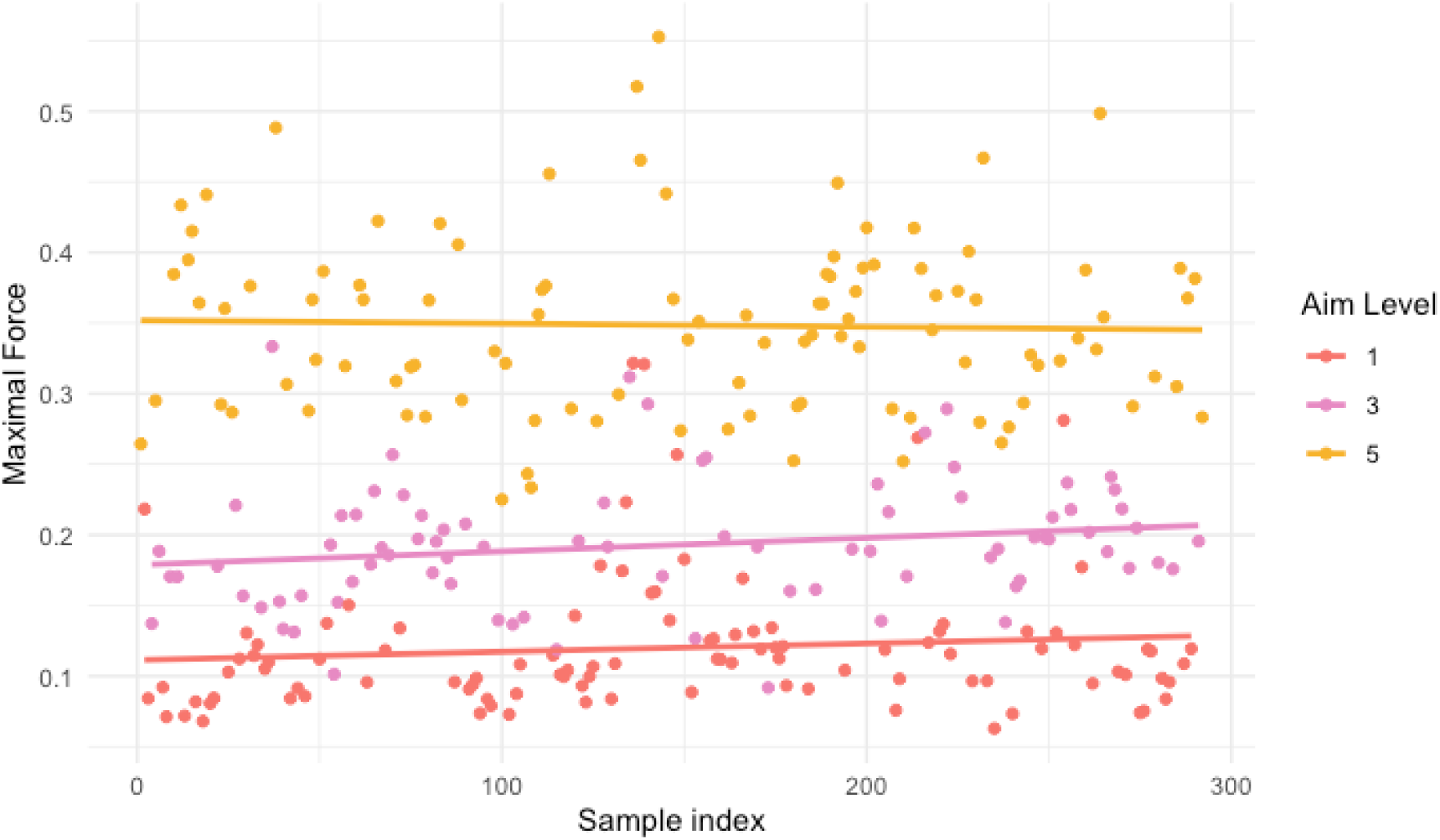
Linear regressions of all three conditions for a representative participant. Data is normalized per participant’s MVC but not standardized. N = 119.

**Figure 7.**
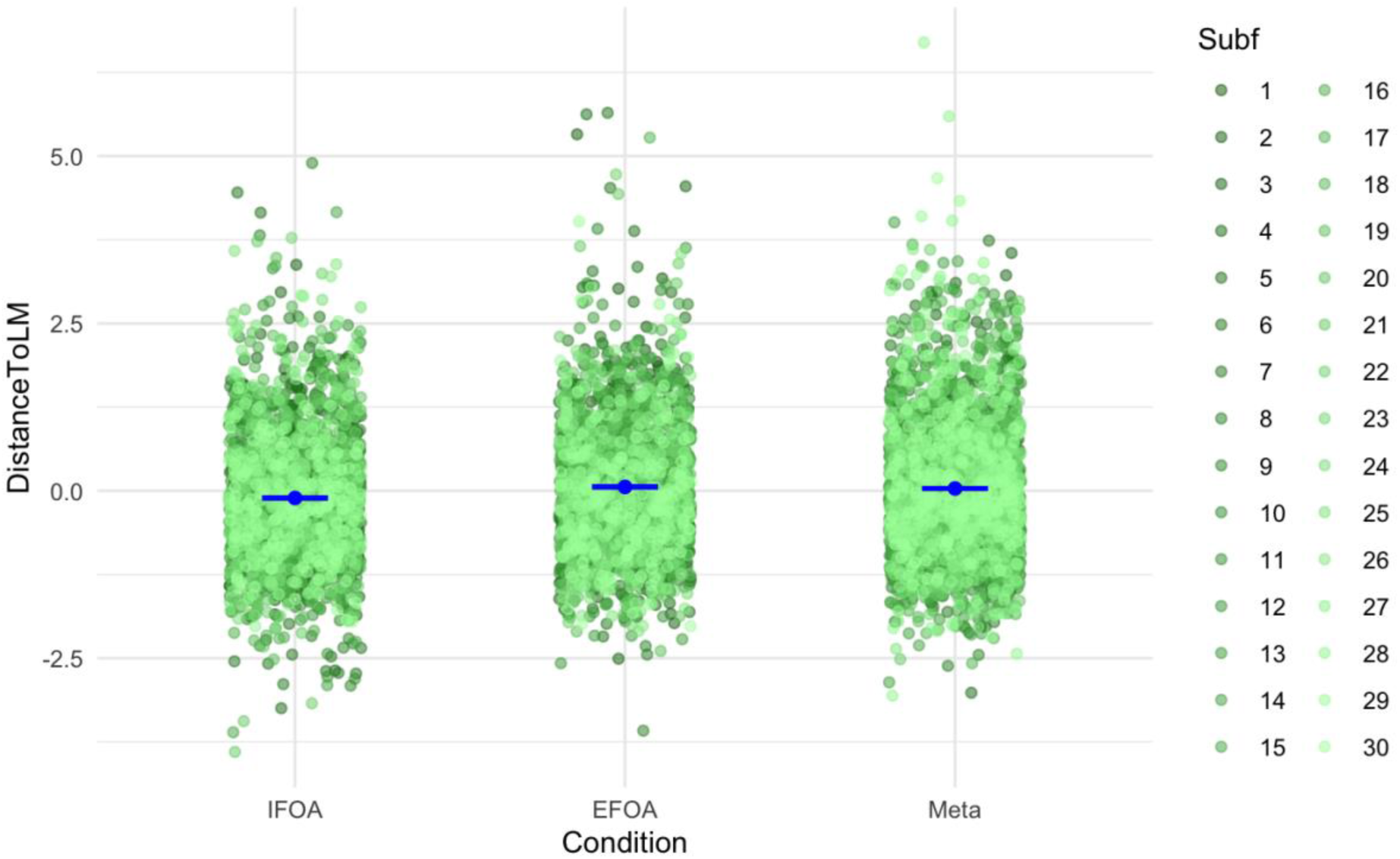
Difference between force and linear regression predicted force for each condition, meta, IFOA, and EFOA, standiardized per subject and aim, colored by participant. 0 Signifies no difference, negative is undershooting and positive is overshooting. Generally, the distance is much smaller than the distance to MVC (fig. 4). N = 8390.

**Table 4.** Left: Estimates of a brm model with distance between force and LM predicted force standardized per subject and aim as the dependent variable and condition and aim as the independent variables. Right: Hypothesis tests whether there is difference in the distance between force and lm predicted force between conditions. The distance to lm is lower for IFOA than both other conditions. This indicates that force is less variable around the internal representation of the force level in IFOA than in meta and EFOA.

|  | Estimate | Est. Error | 95% CI | Hypothesis test | Estimate | Est. Error | 95% CI | Evid. Ratio | Post. Prob |
| --- | --- | --- | --- | --- | --- | --- | --- | --- | --- |
| Intercept | -0.07 | 0.02 | [-0.12; -0.02] | Meta - IFOA > 0 | 0.22 | 0.04 | [0.14; 0.29] | Inf | 1.00 * |
| EFOA | 0.17 | 0.03 | [0.12; 0.22] | EFOA - IFOA > 0 | 0.24 | 0.05 | [0.17; 0.32] | Inf | 1.00 * |
| Meta | 0.15 | 0.02 | [0.10; 0.19] | EFOA - Meta > 0 | 0.03 | 0.02 | [-0.01; 0.06] | 5.51 | 0.85 |
| Level 3 | -0.04 | 0.03 | [-0.09; 0.01] |  |  |  |  |  |  |
| Level 5 | -0.05 | 0.02 | [-0.10; -0.00] |  |  |  |  |  |  |

To test the impact of condition on the distance from actual force to the linear regression predicted force (distance to LM), we first standardized the difference to MVC by aim to align variance. Then we estimated a Bayesian linear mixed model containing the DistanceToLM as the dependent variable and condition and subject as the independent variables (Fig. 7, Table 4). We used Bayesian hypothesis testing to evaluate whether DistanceToLM differs between conditions (Fig. 7, Table 4). We found that distance to LM was larger in the meta condition and EFOA condition compared to the IFOA condition (Est. (IFOA) = -0.07, Est. Error = 0.02 [-0.12; -0.02], Est. (EFOA) = 0.17, Est. Error = 0.03 [0.12; 0.22], Est. (meta) = 0.15, Est. Error = 0.02 [-0.10; 0.19]). Hypothesis tests showed that meta estimates are larger than IFOA estimates of distance to LM with infinite evidence ratio and Post. Prob = 1.00, and that EFOA estimates are larger than IFOA estimates of distance to LM with infinite evidence ratio Post. Prob = 1.00 *. This indicates that in the IFOA condition, the variability in force is smaller around participant’s internal representation of the aim levels as estimated by the lm.

To test whether the distance to LM is explanatory of the MCJ in the meta condition, we estimated a Bayesian linear regression model with the difference between force and MVC as the dependent variable and metacognitive judgement and subject as the independent variables (Fig. 8, Table 5). We performed Bayesian hypothesis tests to evaluate whether the estimates for the force levels differ between MCJ levels (Fig. 8, Table 5).

**Table 5.**
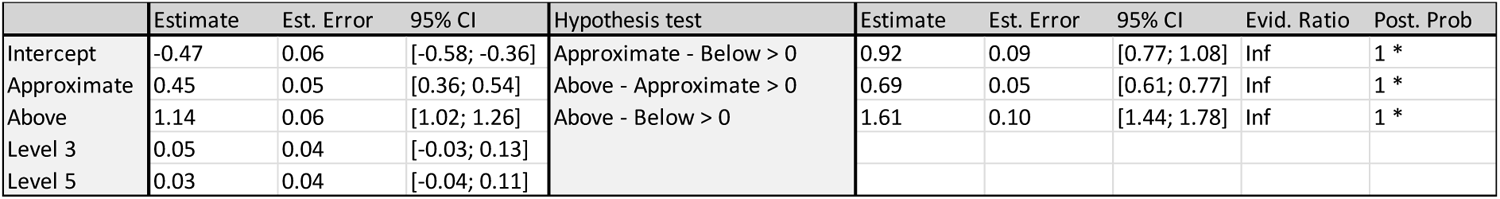
Distance to LM estimates between MCJs, distance to LM standardized per subject and aim.

**Figure 8.**
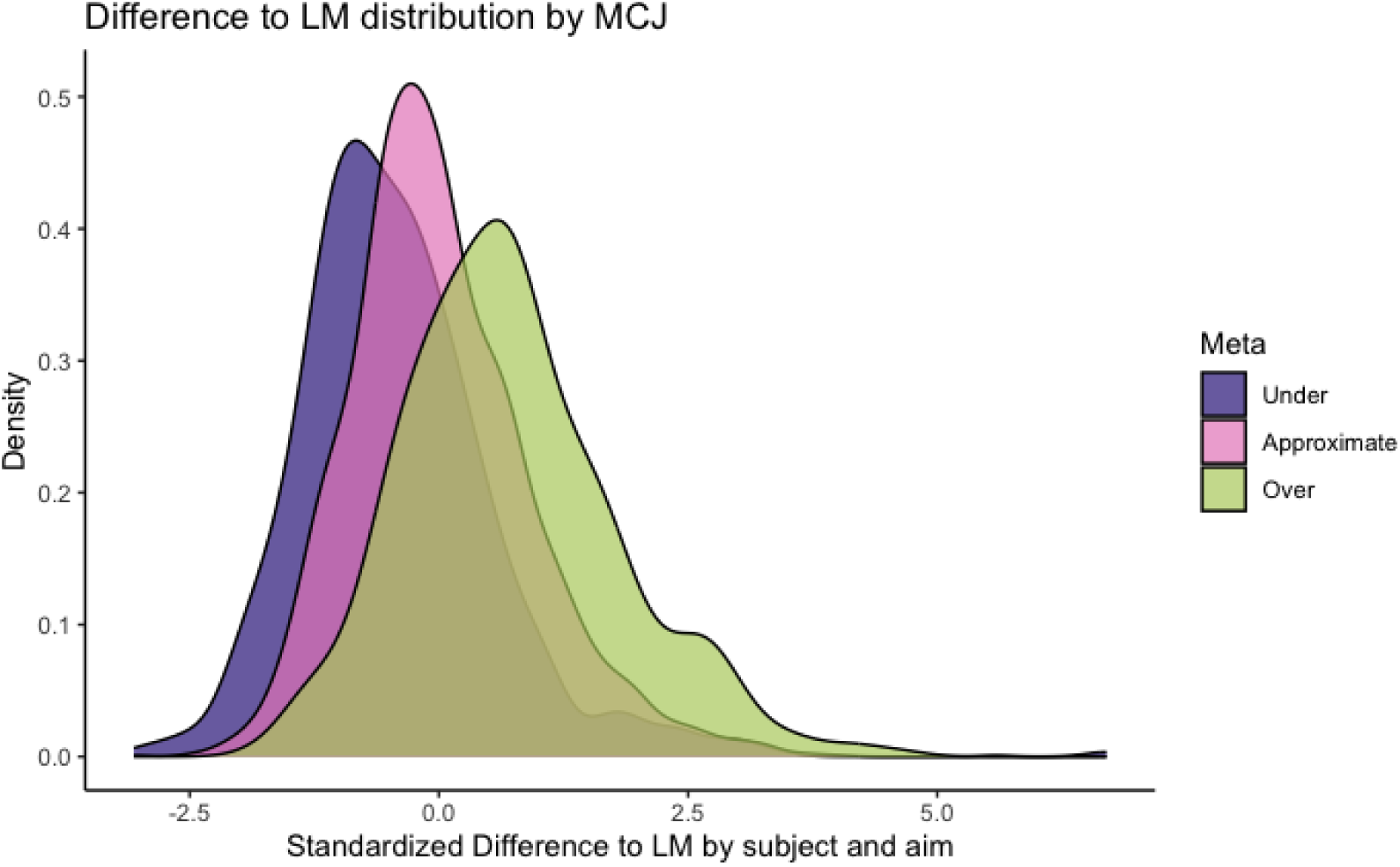
Standardized distance to LM by participant and by aim, distributed by MCJ. N= 3510.

We find that the estimates of the distance to LM increase with MCJ, (Est.(below) = -0.47, Est. Error = 0.06 [-0.58; -0.36], Est. (approximate) = 0.45, Est. Error = 0.05 [0.36; 0.54], Est (above) = 1.14, Est. Error = 0.06 [1.02; 1.26]). Hypothesis tests showed that these effects were significant with infinite evidence ratio and a posterior probability of 1 (Table 4, Fig. 9). Together, these results support the findings from the difference to %MVC; estimates of difference between force and the linear regression predicted force increases with MCJ follows distance to LM, indicating that participants correctly assign the MCJs.

**Fig. 9.**
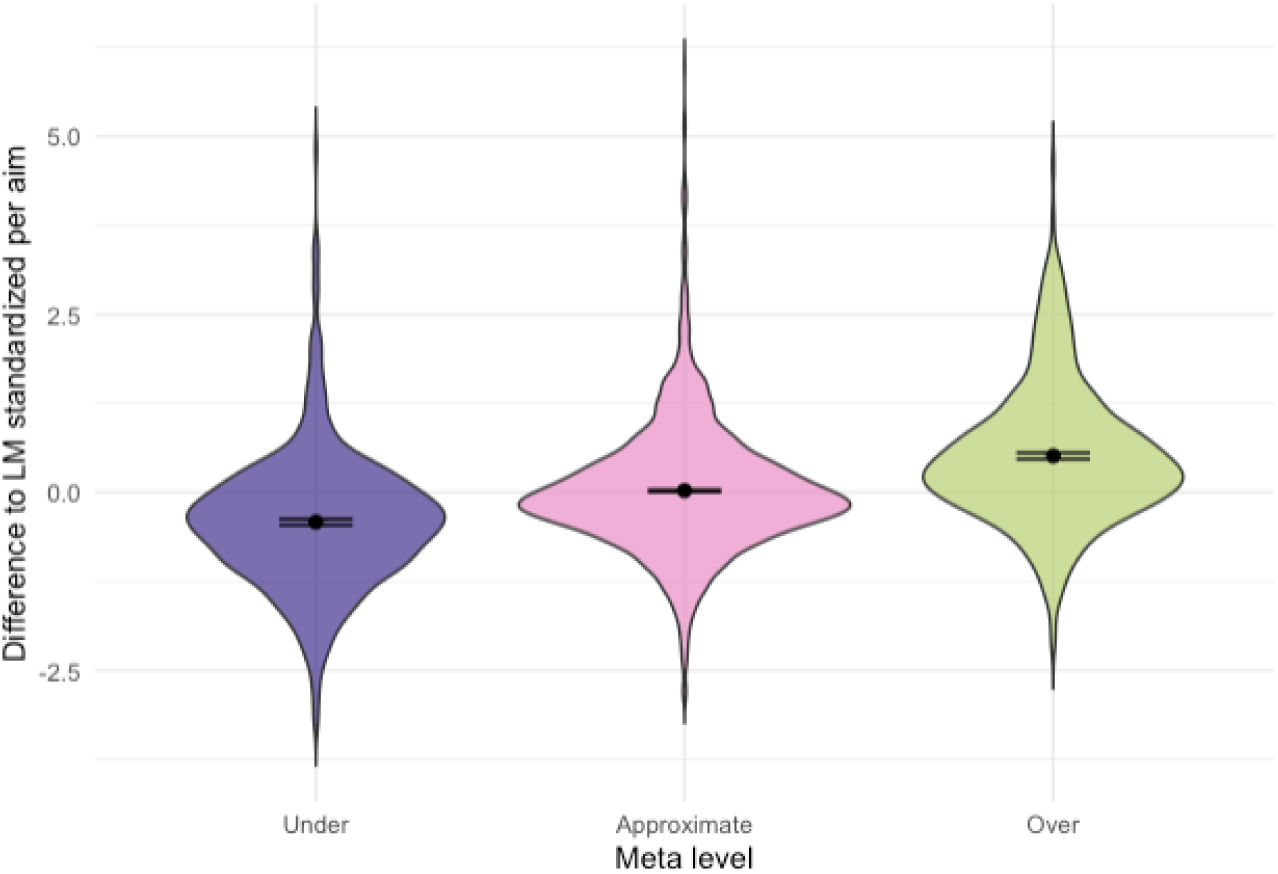
Distance to LM distributed by metacognitive judgment standardized by aim. N = 3510.

As we do not control for first-order behavior (in our case the variance of force produced), we instead tested the relationship between metacognitive access and variance in motor performance in three ways. First, we tested the relationship between percentage of correct answers and variance in motor performance (Fig. 10a). Participants differed in the amount they assigned ‘off’ (below or above) compared to ‘on’ (approximate). To account for this, we multiplied the percentage correct answers by a ratio of off/all assignments, and tested whether this measure was correlated with motor variance (Fig. 10b). Lastly, we computed a ratio of variance in all measures divided by variance in ‘on’ measures, and tested whether this was correlated with motor variance (Fig. 10c). For all measures we found no correlation to motor variance (Table 6). Notice, that for some participants, the variance in ‘off’ is not larger than ‘on’ (they have an estimate around 1), indicating that these participants do not have metacognitive access to their movements.

**Figure 10.**
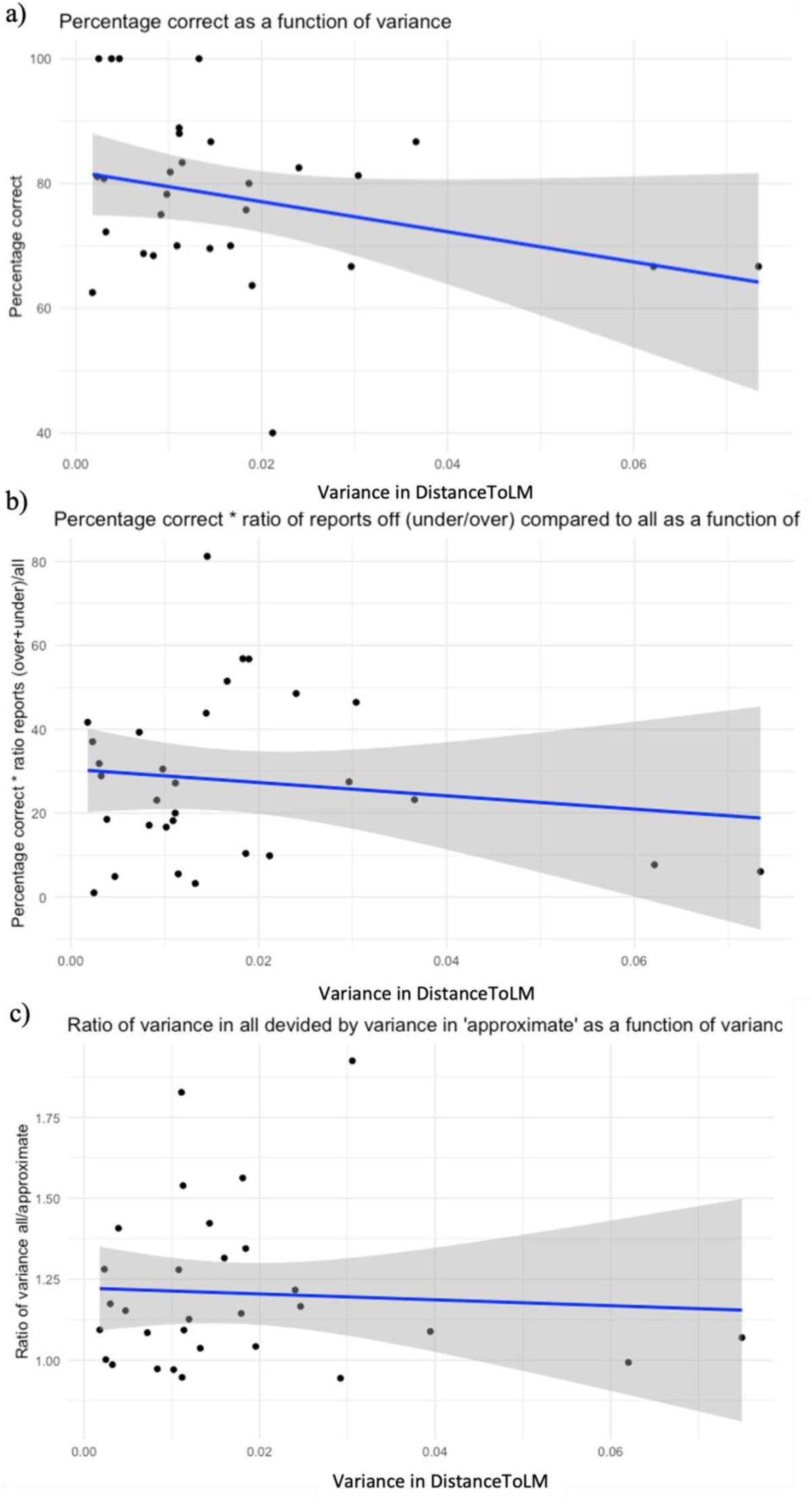
Correlations between metacognitive ability and variance in DistanceToLM. A) Correlation between percentage of correct answers and variance in DistanceToLM. B) Correlation between percentage of correct answers multiplied with off/all ratio and variance in DistanceToLM. C) Correlation of the ratio of variance in all measures divided by variance in ‘on’ measures and variance in DistanceToLM. N = 30.

**Table 6.** Bayesian hypothesis tests of correlations between metacognitive ability and variance in DistanceToLM. Correlation between percentage of correct answers and variance in DistanceToLM. Correlation between percentage of correct answers multiplied with off/all ratio and variance in DistanceToLM. Correlation of the ratio of variance in all measures divided by variance in ‘on’ measures and variance in DistanceToLM.

|  |  | Estimate | Est. Error | 95% CI | Hypothesis test | Estimate | Est. Error | 95% CI | Evid. Ratio | Post. Prob |
| --- | --- | --- | --- | --- | --- | --- | --- | --- | --- | --- |
| Percent correct | Intercept | 77.9 | 2.47 | [73.0; 82.8] | Variance < 0 | 0.01 | 1 | [-1.62; 1.67] | 0.99 | 0.5 |
|  | Variance | 0.01 | 1.00 | [-1.97; 1.98] | Variance > 0 | 0.01 | 1 | [-1.62; 1.67] | 1.01 | 0.5 |
| Percent correct * ratio respons | Intercept | 18.5 | 1.88 | [20.8; 34.8] | Variance < 0 | -0.01 | 1.01 | [-1.69; 1.64] | 1.01 | 0.5 |
|  | Variance | -0.02 | 0.98 | [-1.94; 1.89] | Variance > 0 | -0.01 | 1.01 | [-1.69; 1.64] | 0.99 | 0.5 |
| Variance ratio | Intercept | 1.21 | 0.05 | [1.11; 1.31] | Variance < 0 | -0.12 | 0.98 | [-1.73; 1.47] | 1.23 | 0.55 |
|  | Variance | -0.11 | 0.94 | [-1.96; 1.76] | Variance > 0 | -0.12 | 0.98 | [-1.73; 1.47] | 0.81 | 0.45 |

## Discussion

### Main findings

The main aim of this study was to investigate our ability to consciously monitor own motor signals. First, we tested whether participants have metacognitive access to variability in own force between trials in an index finger movement task with no visual feedback. We found that participants can correctly assign metacognitive judgements (MCJs) to their own force. This suggests that participants do indeed have metacognitive access to their sensorimotor signals. Second, we tested whether there is a difference in inter-trial force variance between different foci of attention without visual feedback. We found that when participants attend to their internal perception of the movement, the inter-trial force variance is decreased compared to when they focus on imagined external feedback or on evaluating the force level in the meta condition. This indicates that internal focus of attention (IFOA) is superior to other foci of attention in reducing inter-trial force variance. Third, we tested the relationship between metacognitive access to sensorimotor signal and variance in motor performance, and found no correlation between the two.

### Task validation

We found that on a group level, participants can consistently produce three distinct force levels without visual feedback in all conditions, IFOA, EFOA, and meta. Force measures increase with aim level, and force distributions per aim are different with infinite evidence ratio and a posterior probability of 1. Importantly, with these behavioral results we replicate findings from a prior study (Brandt et al., n.d.). Participants performed comparably across conditions in producing different force distributions for each aim level, although estimates of aim level 3 were higher in the meta condition compared to the EFOA condition.

### Metacognitive judgments

To evaluate whether participants correctly assign MCJs in the meta condition, first, we tested whether MCJs were informative of the distance to the aim force level, which was quantified as a percentage of MVC (10%, 30%, and 50% of own MVC). To further evaluate the accuracy of MCJs, we investigated whether MCJs could reliably indicate the discrepancy between participants’ actual force output and their internal representation of the desired force level (distance to LM). This internal representation was determined by predicting the force for each trial using a linear regression model. This model was fitted on the force data from all other trials within the same block for each specified aim level. For both distance to MVC and distance to LM, hypothesis tests showed that the force estimate systematically increased with each MCJ level from ‘below’ to ‘approximate’ to ‘above’. This pattern suggests that, on a group-level, participants were effectively able to correctly assign MCJs. The results for distance to MVC demonstrate not that participants were overshooting or undershooting the specified %MVC, but rather that the estimates of the force exerted linearly aligned with their MCJs. The results for the distance to LM indicate that participants can distinguish between when they are below, approximate, or above their internal representation of the force level. The difference to MVC is generally larger than the difference to LM. This is not surprising, it indicates that participants hold an internal representation of the levels that they center their force levels around, which differs from the externally imposed percentage MVC. Together, these results confirm our hypothesis that participants do in fact have (metacognitive) access to variability in their own force in an index finger motor task without visual feedback. Correctly assigning metacognitive judgments indicates that participants have access to an error signal on which to base their judgment. This error signal may arise either between intention and perception or between perception and motor memory.

We do not control for first-order behavior (the variance of force produced). For a participant with low inter-trial force variance, it will be a more difficult starting point to assign metacognitive judgments, when the variance to assign it on is smaller than for someone with high inter-trial force variance. Since we do not have a forced-choice task but rely on participants own movements, it was not possible to use a staircasing method to control for this first-order behavior. Therefore, we tested the relationship between metacognitive ability, measured in three different ways, and variance in the motor task. For all three tests we found no significant relationship between metacognitive access and motor variance. This indicates that motor performance does not determine the ability to consciously monitor one’s own movements. In the last test we compared a ratio of the variance in all force measures and the variance in trials in which participants assigned the MCJ ‘approximate’, to variance in all trials. We did this to test whether the variance in the trials in which participants judged the force to be approximate was smaller than the variance in all trials, including the trials in which they judged the force to be ‘below’ or ‘above’. This ratio is used as a measure of metacognitive access, and we tested whether this correlated with the variance in force measures. We found no correlation, but the results suggest that participants differ in their metacognitive ability. Some participants have a ratio around 1, indicating no metacognitive access to sensorimotor signals, whereas other participants have a ratio of 2, indicating high metacognitive access to sensorimotor signals.

Our results bring new perspectives to earlier studies. They challenge the conclusion from visuomotor discrepancy studies such as the one by Fourneret and Jeannerod (1998), which suggested that our access to sensorimotor information is limited (Fourneret and Jeannerod, 1998). Contrary to these conclusions, our results suggest that some participants can in fact consciously access detailed sensorimotor information. Similarly, Charles et al. (2020) observed that while sensorimotor information did not enhance metacognitive efficiency, it did increase accuracy in a task requiring participants to discern whether a probe appeared in front of or behind their hand (Charles et al., 2020). Their results suggest that sensorimotor information improves accuracy in conscious monitoring of the movement when the sensorimotor information is not in discrepancy with the visual information.

The divergent findings between, on the one hand, the visuomotor discrepancy studies and, on the other hand, our study and Charles et al. (2020) may stem from differences in the mechanisms of sensorimotor integration. It appears that visuomotor discrepancies lead to impaired conscious monitoring of one’s own sensorimotor signals. However, in the absence of such discrepancies, sensorimotor information is accessible to some participants and enhances accuracy (Charles et al., 2020). This suggests that sensory integration plays a critical role in perception of movement.

Kasuga et al (2022) suggest a two-stage process for integration of sensory information. In an initial subcortical stage, information from each sensory modality is processed separately, allowing fast error-correction based on only proprioceptive feedback, whereas in the second cortical stage, different sensory modalities are integrated (Kasuga et al., 2022). In the second stage, when information from various modalities is integrated into a unified experience, visual information dominates. This could be due to lower variance in the visual information. Ernst and Banks et al. (2002) suggest that the visual information dominates the percept when the variance of the visual estimation is lower than the haptic estimation (Ernst and Banks, 2002). Taken together, our findings suggest that individuals are capable of consciously monitoring deviations in sensorimotor signals. We suggest that previous results can be attributed to the dominance of visual information, which significantly influences the perception of sensorimotor information. It appears that when visual and sensorimotor information conflict, the low variance and the reliability of visual signals tends to overshadow sensorimotor information. However, in the absence of such discrepancies, agents can consciously access and utilize sensorimotor information effectively. This highlights the complex interplay between different sensory modalities in shaping our percept and underlines the importance of sensorimotor signals. We argue, therefore, that it is important with studies focusing exclusively on movement when investigating the awareness of motor signals.

### Focus of attention

Distance to MVC did not differ between the IFOA, EFOA, and meta conditions, suggesting that participants are equally capable of replicating an externally defined force level across these three different attentional focuses, even without visual feedback. Interestingly, we observed a difference in distance to LM between conditions. We found that the distance was smaller in the IFOA condition compared to both EFOA and meta conditions, indicating that the variance within each force level is smaller when participants hold an IFOA.

This finding is notable given that an EFOA is often associated with enhanced motor performance (Lohse et al., 2014b; Wulf, 2013). However, studies with this finding typically include tasks with an external goal in which visual information is also needed to complete the task. In these cases, the EFOA condition benefits by the visual information to a higher degree than the IFOA condition. In contrast, the IFOA condition compels participants to divert their attention away from externally relevant signals, thereby potentially skewing the advantage towards EFOA.

Lohse et al. (2014) proposed that an EFOA reduces variance in the outcomes we attend to. For instance, in their dart throwing experiment, EFOA decreased endpoint variability in dart placement, whereas having an IFOA on elbow movements decreased variance in elbow movements (Lohse et al., 2014a). Our findings align with this theory, suggesting that the effect of focus is task-dependent: EFOA enhances performance when the goal is external, whereas IFOA proves beneficial when the aim is to minimize movement variance.

Under IFOA, participants are paying attention to the available information in the motor system. Paying attention to this information improves the movement precision. This indicates that focusing on internal cues likely enhances the use of internal information from the motor system. Taking an optimal control theory point of view, the internal information used could be both the intention, afferent information (motor command) and efferent information (proprioceptive feedback). Our results suggest that taking an IFOA facilitates better integration and utilization of this information. This indicates, that not only is it possible to access sensorimotor information, it is also possible to use conscious attention to improve our movements.

### Limitations

Some participants significantly exceeded the intended % MVC during the task, with force estimates approaching or even surpassing their MVC at aim 5, despite being targeted at only 50%. This discrepancy could stem from variations in how participants apply force to the transducer during MVC measurements compared to task performance. The MVC for each participant is determined by the highest force recorded during one of two three-second presses, separated by a 90-second rest interval. However, during the task, participants are instructed to execute shorter presses. Shorter presses can allow for more forceful exertions, potentially enabling participants to exceed the force levels recorded during their MVC determination. For other participants, estimates of the force levels are clearly lower than the percentage MVC they should have performed. The divergence from the MVC scale explains that the range in the difference to MVC is large. We find that this is a necessary consequence of a setup in which we withdraw external feedback, and this is why we extend the analysis not just focusing on an externally imposed scale that is relative to each participant’s MVC, but also on an internal representation of the force levels calculated through linear regressions, distance to LM. For two of the participants, the linear regressions across different force levels overlapped. This overlap could be attributed to one of two possibilities: Either these participants were unable to perform distinct force levels, or they had difficulty distinguishing between the tone pitches that signaled the required force level. Since the participants did not report any issues with discerning the tones, we chose to include them in our analysis.

That the internal representation of force level changes linearly over the course of the experiment is an assumption we are making. The internal representation could possibly change non-linearly.

### Conclusions and further perspectives

On a group-level, participants were able to correctly monitor the force variations in their own movements without visual feedback. Some participants showed higher variance in the ‘off’ trials compared to the ‘on’ trials, indicating that they had metacognitive access to a sensorimotor error signal. Other participants did not show this metacognitive access. Knowing whether we have perceptual access to the deviation/error in our movement is the first step toward investigating the functional role of perception.

Further, our results indicate that during IFOA participants exhibit lower variance in force per aim level. This indicates that IFOA decreases variance in motor performance compared to EFOA in this task, which suggests that IFOA might be beneficial when decreasing variance in movements that do not rely on external feedback.

The current study extends existing literature by focusing exclusively on MCJs of motor performance, rather than sensory integration with visual stimuli. With this study we find that a subset of participants has access to sensorimotor error signals. Although motor control processes can and often do happen without explicit monitoring (Blakemore et al., 2002), it seems that a subgroup can monitor it if they intend to focus on it.

Future studies could explore the potential of using conscious monitoring of sensorimotor signals to enhance motor learning in the absence of external feedback. Could the MCJ be used as feedback to optimize motor output in subsequent attempts? Also, investigating the neural correlates of error detection that underpin MCJs could illuminate the brain mechanisms involved, and potentially shed light into which sensorimotor signals the MCJs rely on.

## Additional information

### Data availability

The data supporting the findings in this study are available upon reasonable request from the corresponding author IMB. The data are not publicly available yet since the data are currently pseudo-anonymized. We will fully anonymize the data and make it available with Open Access at Open Science Foundation in case of manuscript acceptance.

### Code availability

The code supporting the findings in this study is available upon reasonable request from the corresponding author IMB. We will make it available with Open Access at Open Science Foundation in case of manuscript acceptance.

## Author contributions

Ida Marie Brandt: Conceptualized and designed the study, organized data collection, performed data analysis, discussed results, and wrote the original manuscript draft of the paper and additional editorial corrections.

Sofie Maria Bjerregaard: Performed data collection. David Bjerregaard Jensen: Performed data collection.

Markus Grenaa Giessing: Performed data collection and edited the manuscript.

Thor Grünbaum: Conceptualized the study, provided feedback on experimental setup, discussed results, edited the manuscript.

Mark Schram Christensen: Obtained funding, conceptualized the study, discussed results, edited the manuscript.

## Supplementary material

**Supplementary table 1.**
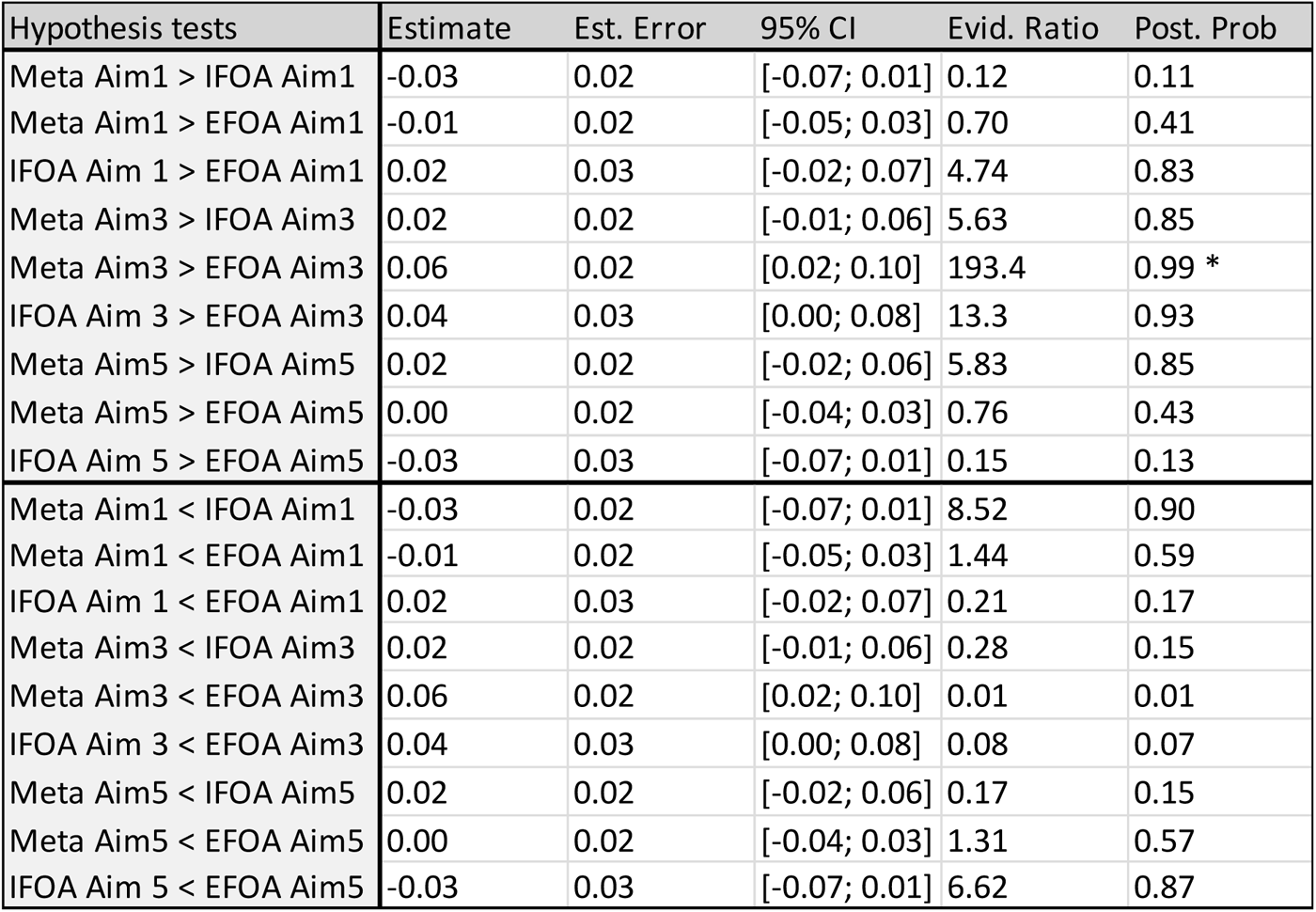
Bayesian hypothesis tests of the interaction between condition and aim level. The aim levels do not differ between conditions, except for the Aim 3 distribution, which is higher in the meta condition compared to the external focus of attention condition. EFOA: External focus of attention, IFOA: Internal focus of attention.

| Hypothesis tests | Estimate | Est. Error | 95% CI | Evid. Ratio | Post. Prob |
| --- | --- | --- | --- | --- | --- |
| Meta Aim1 > IFOA Aim1 | -0.03 | 0.02 | [-0.07; 0.01] | 0.12 | 0.11 |
| Meta Aim1 > EFOA Aim1 | -0.01 | 0.02 | [-0.05; 0.03] | 0.70 | 0.41 |
| IFOA Aim 1 > EFOA Aim1 | 0.02 | 0.03 | [-0.02; 0.07] | 4.74 | 0.83 |
| Meta Aim3 > IFOA Aim3 | 0.02 | 0.02 | [-0.01; 0.06] | 5.63 | 0.85 |
| Meta Aim3 > EFOA Aim3 | 0.06 | 0.02 | [0.02; 0.10] | 193.4 | 0.99 * |
| IFOA Aim 3 > EFOA Aim3 | 0.04 | 0.03 | [0.00; 0.08] | 13.3 | 0.93 |
| Meta Aim5 > IFOA Aim5 | 0.02 | 0.02 | [-0.02; 0.06] | 5.83 | 0.85 |
| Meta Aim5 > EFOA Aim5 | 0.00 | 0.02 | [-0.04; 0.03] | 0.76 | 0.43 |
| IFOA Aim 5 > EFOA Aim5 | -0.03 | 0.03 | [-0.07; 0.01] | 0.15 | 0.13 |
| Meta Aim1 < IFOA Aim1 | -0.03 | 0.02 | [-0.07; 0.01] | 8.52 | 0.90 |
| Meta Aim1 < EFOA Aim1 | -0.01 | 0.02 | [-0.05; 0.03] | 1.44 | 0.59 |
| IFOA Aim 1 < EFOA Aim1 | 0.02 | 0.03 | [-0.02; 0.07] | 0.21 | 0.17 |
| Meta Aim3 < IFOA Aim3 | 0.02 | 0.02 | [-0.01; 0.06] | 0.28 | 0.15 |
| Meta Aim3 < EFOA Aim3 | 0.06 | 0.02 | [0.02; 0.10] | 0.01 | 0.01 |
| IFOA Aim 3 < EFOA Aim3 | 0.04 | 0.03 | [0.00; 0.08] | 0.08 | 0.07 |
| Meta Aim5 < IFOA Aim5 | 0.02 | 0.02 | [-0.02; 0.06] | 0.17 | 0.15 |
| Meta Aim5 < EFOA Aim5 | 0.00 | 0.02 | [-0.04; 0.03] | 1.31 | 0.57 |
| IFOA Aim 5 < EFOA Aim5 | -0.03 | 0.03 | [-0.07; 0.01] | 6.62 | 0.87 |

